# Geometric coupling of helicoidal ramps and curvature-inducing proteins in organelle membranes

**DOI:** 10.1101/635466

**Authors:** Morgan Chabanon, Padmini Rangamani

## Abstract

Cellular membranes display an incredibly diverse range of shapes, both in the plasma membrane and at membrane bound organelles. These morphologies are intricately related to cellular functions, enabling and regulating fundamental membrane processes. However, the biophysical mechanisms at the origin of these complex geometries are not fully understood from the standpoint of membrane-protein coupling. In this work, we focused on a minimal model of helicoidal ramps representative of specialized endoplasmic reticulum compartments. Given a helicoidal membrane geometry, we asked what is the distribution of spontaneous curvature required to maintain this shape at mechanical equilibrium? Based on the Helfrich energy of elastic membranes with spontaneous curvature, we derived the shape equation for minimal surfaces, and applied it to helicoids. We showed the existence of switches in the sign of the spontaneous curvature associated with geometric variations of the membrane structures. Furthermore, for a prescribed gradient of spontaneous curvature along the exterior boundaries, we identified configurations of the helicoidal ramps that are confined between two infinitely large energy barriers. Overall our results suggest possible mechanisms for geometric control of helicoidal ramps in membrane organelles based on curvature-inducing proteins.

## Introduction

Recent advances in microscopy, segmentation, and reconstruction methods have enabled us to visualize the plasma membrane and intracellular organelle membranes with a previously unforeseen level of spatial resolution [1–4]. These observations demonstrate the stunning complexity of membrane architecture within cells. Remarkably, despite this diversity, some common geometrical features are found across organelles and functions [5]. For instance budding/necking processes occur in the plasma membrane through a variety of endocytic and exocytic mechanisms (e.g., clathrin mediated [6–8], ESCRT [9–11], antimicrobial peptides [12, 13]), but also in the endoplasmic reticulum (ER) and Golgi in anterograde and retrograde trafficking [14, 15]. Similarly membrane tubes are present in the ER membrane network [1, 16–18], mitochondria [19–21], and in the form of cellular nanotubes [22–24]. Common to these seemingly disparate membrane structures is the catenoid-like geometry appearing at the neck of buds and the base of tubes [25]. Another striking example is the recently identified helicoidal structure connecting membrane sheets in the peripheral ER of neural and secretory cells [1, 26] as well as in the spine apparatus present in dendritic spines [4] (see Fig. 1). The molecular and biophysical mechanisms by which these helicoidal membrane geometries are produced and maintained remain under active investigation. However, the conserved features of membrane shapes across different cellular locations and biological functions suggest the possible existence of common design principles relating lipid membrane geometry, composition, and function [5].

**Figure 1:**
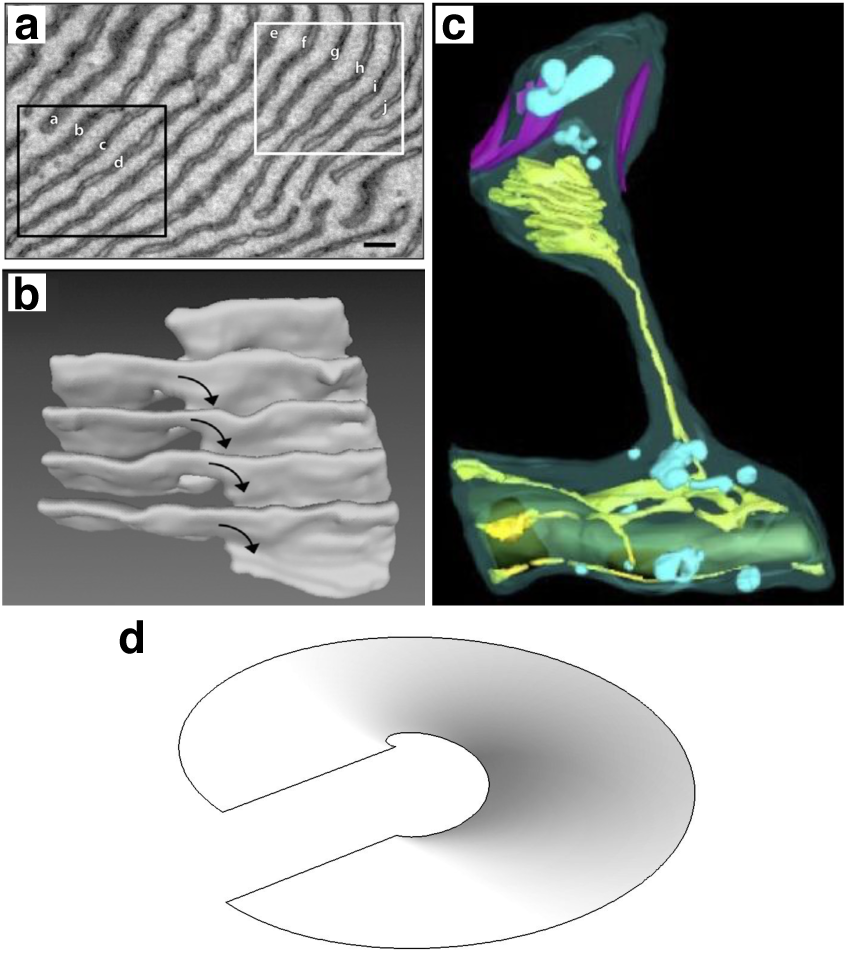
Helicoidal membrane structures in the endoplasmic reticulum (ER). (a) Electron microscopy cross section of the ER in saliva secretory cell, (adapted from [26]). (b) Three-dimensional reconstruction of the black square in (a) showing a helicoidal ramp structure. (c) Spine apparatus in dendritic spine (adapted from [4]). (d) Geometry of a helicoidal ramp surface.

We previously proposed a minimal model of membrane-protein interactions in catenoid-like necks [25], where our main finding was that at least two distinct curvature-inducing mechanisms are required to constrain a catenoid membrane neck below a critical radius. This has broad implications in ESCRT-mediated budding [9–11], yeast mitochondrial fission by dynamin [27], as well as yeast budding by septin filaments [28]. The fact that helicoids and catenoids belong to the same family of minimal surfaces – mathematical objects that minimize their surface energy by having a zero mean curvature and a negative Gaussian curvature everywhere – suggests that some of the insights gained from catenoids can be extended to helicoids. Following the idea that simple geometric arguments can provide insight into fundamental mechanisms underlying cellular organization [29], we sought to understand the relationship between curvature-inducing proteins and the geometry of intracellular lipid structures, in particular on helicoids representative of ER ramps (Fig. 1).

In a recent study that revealed the architecture of ER ramps to have helicoidal shapes, Terasaki *et al.* [26] proposed a physical model of the helicoidal membrane structures. They considered the mid-plane surface between the two ER bilayers – assumed to be at a constant distance from each other – bordered by an edge line representing the curved axis of the hemicylinder joining the two bilayers. They minimized the bending energy and the effective line energy of such a system. For an edge line having a negative spontaneous curvature [26, 30], they found that the structures satisfying the minimal energy are hollow helicoids such as the one shown in Fig. 1d. Extending this model to account for a mixture of membrane-proteins inducing either constant or negative edge curvatures, Shemesh *et al.* [30] numerically recapitulated various membranes structures present in the ER. In particular, they found that helicoid geometries are favored when the total amount of proteins is low with predominantly negative curvature-inducing proteins at the edges. While these studies lay the foundation of ER structure mechanics, they have focused on the influence of curvature-inducing proteins at the edges only [31], neglecting any potential coupling between curvature-inducing proteins at the sheet surface and the helicoid geometry.

Here we sought to understand the coupling between the distribution of curvature-inducing proteins and the geometry of a helicoidal membrane. We used *spontaneous curvature* as the mathematical catch-all for the curvature induced in the membrane by various molecular mechanisms. Using helicoids as geometric inputs, we computed the distribution of spontaneous curvature that satisfies the mechanical equilibrium of membrane bending [32]. This allowed us to map a geometric state-space for helicoidal membranes in which the bending energy is constrained by protein-induced curvature distribution.

## Model development

### Model assumptions

The main assumptions underlying the model are:

a. The lipid membrane is treated as a continuous thin elastic film of negligible thickness [32]. This assumption relies on the length-scale separation between the membrane thickness (∼ 10^−9^ m) and typical radius of curvature of interest (∼ 10^−8^ − 10^−6^ m).
b. Based on the same length-scale separation, the net effect of compositional asymmetry across the lipid bilayer is accounted for by a “preferred” membrane curvature called *spontaneous curvature* (*C*). This can be induced by several molecular mechanisms, including asymmetric protein insertion in the bilayer, protein scaffolding, clustering, steric pressure, or lipid composition asymmetry [5, 33]. The spontaneous curvature is defined as a penalty for the mean curvature in the Helfrich energy [32] (see Eq. 1).
c. The system is at mechanical equilibrium. Any membrane deformation is much slower than the rate at which the membrane energy is dissipated. Given the coupling between spontaneous curvature and membrane geometry, this also implies no dynamic rearrangement of the membrane composition.
d. The lipid membrane is incompressible and inextensible. This assumption is supported by the orders of magnitude difference between the bending modulus (≃10^−20^ N m) and the stretching modulus (≃0.2 N/m) of lipid membranes [34], allowing us to neglect area variations. This constraint is implemented through a Lagrangian multiplier on the surface area that has the physical interpretation of membrane tension [35, 36].
e. The lipid membrane is inviscid. Because of the fluid properties of lipid bilayers, we neglect any shear forces in the membrane. This simplification is in accordance with assumption (c) where dissipative processes are assumed to equilibrate instantly. Models accounting for in-plane lipid flow and diffusion can be found for instance in [37–39].
f. The lipid membrane is treated as an elastic manifold whose energy functional is the sum of the Helfrich bending energy [32] and a protein entropic contribution [36, 39–41].
g. The bending and Gaussian moduli are assumed to be homogeneous along the membrane surface.

### Elastic lipid membrane with heterogeneous distribution of spontaneous curvature

The most common model of lipid membranes is based on the Helfrich energy [32], which can be extended to account for the entropic contribution of membrane-bound proteins [40] to the area free-energy functional such as

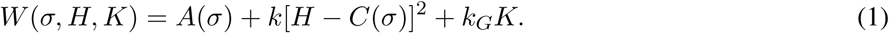

Here *H* is the membrane mean curvature, *K* is the Gaussian curvature, *A*(*σ*) is the contribution of the membrane-bound proteins to the free energy, and *σ* is the surface density of proteins. Furthermore, *k* and *k*_*G*_ are the bending and Gaussian moduli respectively. *C*(*σ*) is the spontaneous (mean) curvature, which is determined by the local membrane composition, say the surface density of a curvature-inducing protein *σ*. While it is certainly possible to propose explicit functions for *A*(*σ*) and *C*(*σ*) of the protein density (see [36, 40] for discussion on *A*(*σ*), and [39, 41] for specific examples) we chose to retain their general form to obtain maximal insight. In the Supplementary Material Section S1, we briefly summarize the derivation of the governing equations of an elastic membrane at mechanical equilibrium to obtain the governing equations (Eqs. S11 and S12 (see [35, 38, 42–45] for detailed derivations).

The shape equation for a lipid membrane with protein-inducing spontaneous curvature is obtained by introducing the free energy density 1 into the normal force balance equation for elastic membranes Eq. S11, resulting in the following partial differential equation

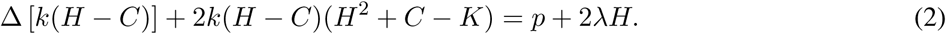

Here *p* is the pressure differential across the membrane and *λ* is a Lagrange multiplier that enforces the inextensibility constraint of the membrane, which is commonly interpreted as the membrane tension [35, 36]. Additionally, Δ(·) is the surface Laplacian. The incompressibility equation for lipid membranes is obtained by introducing Eq. 1 into the tangential force balance equation Eq. S12, resulting in

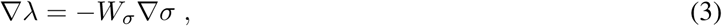

where (·)_*σ*_ = *∂*(·)*/∂σ* is the partial derivative with respect to *σ* and ∇(·) is the surface gradient. *W*_*σ*_ is the chemical potential of the membrane protein, and given Eq. 1, we have *W*_*σ*_ = *A*_*σ*_ − 2*k*(*H* − *C*)*C*_*σ*_.

Eqs. 2 and 3 describe the equilibrium configuration of lipid membrane subject to heterogeneous spontaneous curvature induced by proteins. An additional constraint for imposing the area incompressibility of the membrane requires the lipid velocity field, given by (**u** = *u*^*α*^**a**_*α*_ + *w***n**), to satisfy 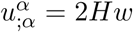 [39]. Provided a lipid velocity field satisfying the above incompressibility condition and suitable boundary conditions exists, the system given by the coupled Eqs. 2 and 3 fully describes the equilibrium configuration of a lipid membrane subject to a static distribution of curvature-inducing proteins. Although models for lipid flow within biological membranes have been proposed [35, 36, 38, 46], such an investigation is out of the scope of this study.

### Distribution of curvature-inducing proteins on minimal surfaces

The general shape and incompressibility conditions expressed by Eqs. 2 and 3 relate the membrane geometry to its local composition. In this section, we specialize the governing equations to minimal surfaces, a mathematical family of surfaces that include helicoids, catenoids, and triply periodic minimal surfaces. We later apply this model to helicoids as minimal models of ER ramps. Using the fact that minimal surfaces have vanishing mean curvature (*H* = 0 everywhere on the membrane), in the absence of transmembrane pressure the shape equation (Eq. 2) reduces to a variable-coefficient Helmhotz equation for the spontaneous curvature

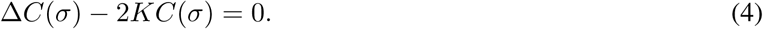

This equation summarizes a key relationship between *C*(*σ*) and *K* in our model. The spontaneous curvature induced by proteins *C*(*σ*) reads the Gaussian curvature of the surface *K*, subject to boundary conditions. Both the energetic contribution of the proteins *A*(*σ*) and the local Lagrange multiplier *λ* are now decoupled from the shape equation for minimal surfaces (Eq. 4), enabling us to solve for those quantities after solving the shape equation. However, any solution of Eq. 4 is restricted to the condition that Eq. 3 is satisfied. For minimal surfaces, this latter equation reduces to

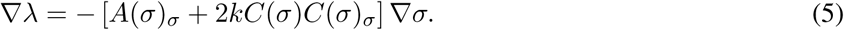

Using the identities ∇*A*(*σ*) = *A*(*σ*)_*σ*_∇*σ* and ∇[*C*(*σ*)^2^] = 2*C*(*σ*)*C*(*σ*)_*σ*_∇*σ* and integrating, we get *λ* as a function of *A*(*σ*) and *C*(*σ*) up to a constant *λ*_0_, such that

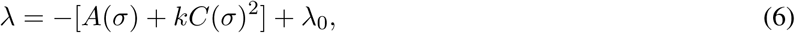

which is the admissibility condition for the Lagrange multiplier field *λ*.

Now that the shape equation is uncoupled from the incompressibility equation, we can solve Eq. 4 for the distribution of spontaneous curvature *C* for a given minimal surface with Gaussian curvature *K*. We complete the system by specifying Dirichlet boundary conditions on the spontaneous curvature *C* = *C*_0_ at the helicoid boundary ramp as indicated in each case [25], while applying a no-flux Neumann boundary condition on other boundaries. The Dirichlet boundary conditions in spontaneous curvature can be understood as the presence of curvature-inducing proteins at the edges of ER ramps and cisternae [30, 47]. On a minimal surface, Eq. 6 not only implies that imposing a Dirichlet condition on the spontaneous curvature is equivalent to imposing a membrane tension at the boundary, but also provides a direct relationship between spontaneous curvature and membrane tension everywhere on the surface.

### Helicoid parametrization

A helicoid surface of diameter *L* and pitch *P* such as the one shown in Fig. 2a can be parametrized as [48, 49]

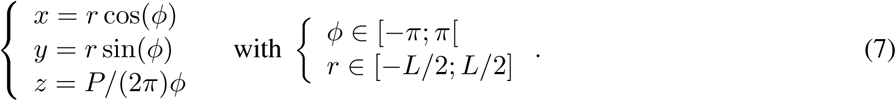

**Figure 2:**
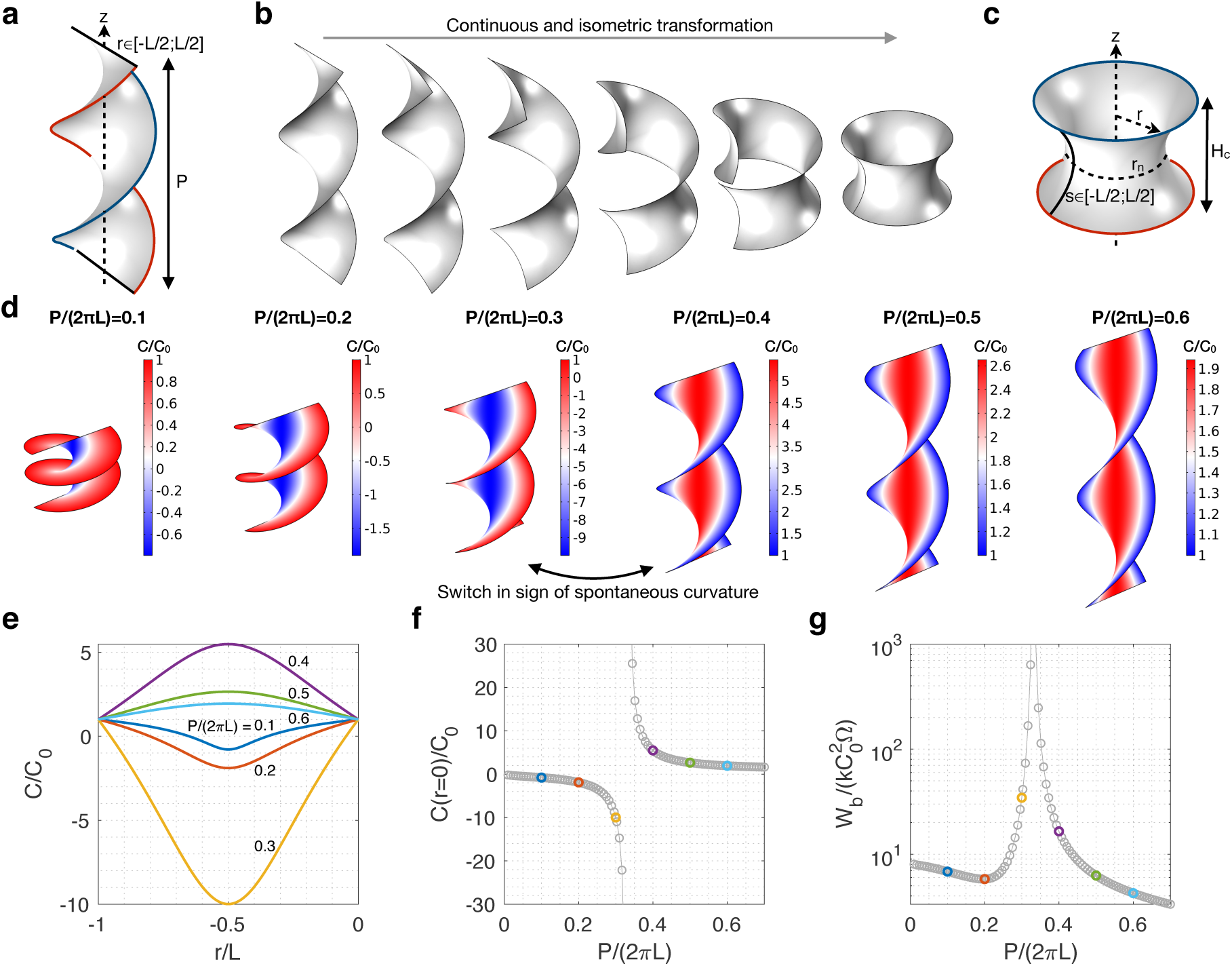
(a-c) Helicoids are continuous and isometric transformations of catenoids. (a) Schematic of a full helicoid and definition of its parameters. (b) Isometric transformation from a helicoid to a catenoid (see Eq. S13). Note that every surface along the transformation is also a minimal surface. (c) Schematic of a catenoid whose boundaries have been transformed from the helicoid shown in (a). The distribution of Gaussian curvature is controlled by the helicoid pitch or equivalently by catenoid neck, through the relation *P* = 2*πr*_*n*_. (d-g) The spontaneous curvature to maintain a helicoid (or a catenoid) shows a switch in sign at a critical pitch (or neck radius). All results here are for boundary conditions set to *C*_0_ = *C*_1_. (d) The computed distribution of spontaneous curvature (normalized by *C*_0_) on full helicoids of varying pitches *P*. (e) Distribution of normalized spontaneous curvature along the radii of the helicoids shown in (d). The spontaneous curvature at the center (*r* = 0) switches sign at a critical pitch between *P/*(2*πL*) = 0.3 and 0.4. (f) Normalized spontaneous curvature at the helicoid center (*r* = 0) as a function of helicoid pitch. Color symbols correspond to the configurations in (e). (g) Corresponding dimensionless bending energy. An energy barrier occurs at a critical helicoid pitch. All results presented for helicoids hold for catenoids providing that the boundary conditions follow the same transformation.

The Gaussian curvature along the radius of such a helicoid is

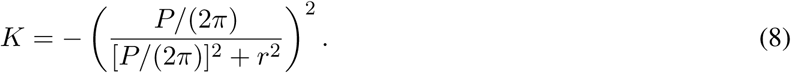

It should be noted that helicoids and catenoids belong to the same associated family of surfaces [49]. So by definition, there exists an isometric transformation from one surface to the other, as illustrated in Fig. 2b (see Supplementary Material Section S2 for details). In particular, the pitch (*P*) of a helicoid is related to the neck radius (*r*_*n*_) of its corresponding catenoid by *r*_*n*_ = *P/*(2*π*). This highlights how, in terms of Gaussian curvature, varying the pitch of a helicoid is equivalent to varying the neck radius of a catenoid. Since the Gaussian curvature enters in the source term of the shape equation for minimal surfaces (Eq.4), the distribution of spontaneous curvature on a helicoid is intrinsically related to its pitch, as a catenoid’s distribution of spontaneous curvature is related to its neck radius [25].

### Dimensionless system

The system has two length scales, a natural length scale *L*, which describes the system size, and an induced length scale, *C*_0_, which arises due to the protein-induced spontaneous curvature. We therefore scale the geometric variables as 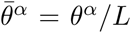 and 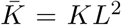, and the spontaneous curvature as 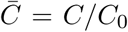. Accordingly, the system can be written in its dimensionless form as

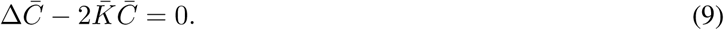

The total membrane bending energy, defined as the integral of the contribution from the curvature to the energy density over the surface (Ω), is given by

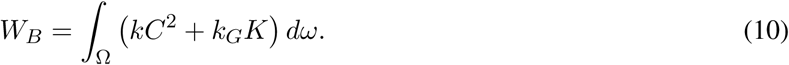

In dimensionless form, the total bending energy is

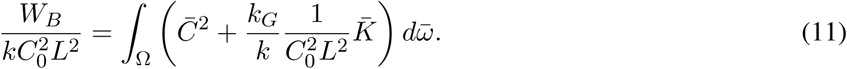

Dividing by the dimensionless area of the surface 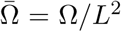, the dimensionless energy per area is 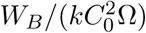. Based on Hu *et al.* [50], we choose the value of the ratio of Gaussian to bending moduli as *k*_*G*_*/k* = −0.9 for the reminder of our analysis.

### Numerical implementation

We solved Eq. 9 subject to either Dirichlet or Neumann boundary conditions using the finite-element solver soft-ware COMSOL Multiphysics^®^ v5.3a (COMSOL AB, Stockholm, Sweden) with the Surface Reaction Module. Parametric studies and quantitative data analysis were conducted in MATLAB R2018b (MathWorks, Inc., Natick, MA, USA) and the COMSOL With MATLAB interfacing module.

## Results

### Distribution of spontaneous curvature on a full helicoid is controlled by the pitch

We first consider helicoids with constant boundary conditions *C*_0_ on both sides of the ramp (Fig. 2a with *C*(±*L/*2) = *C*_0_), for various pitches. This configuration is formally equivalent to solving for the spontaneous curvature on a catenoid for various neck radius as was done in [25], and therefore serves to validate our model (see text associated with Eq. 8 and Supplementary Material Section S2). The resulting distribution of spontaneous curvature is presented as color maps in Fig. 2d and along the helicoid arclength in Fig. 2e. We find that although the maximum intensity of spontaneous curvature is always at the center of the helicoid, its value depends on the pitch. Furthermore, the spontaneous curvature exhibits switches in sign at a critical pitch between *P/*(2*πL*) =0.3 and 0.4. For *P/*(2*πL*) < 0.3, the normalized spontaneous curvature *C/C*_0_ is positive on the outer edge of the helicoid and negative towards the center. For *P/*(2*πL*) > 0.4, *C/C*_0_ at the center switches sign and becomes positive. This switch is better seen in Fig. 2f, where the normalized spontaneous curvature at the center of the helicoid *C*(*r* = 0)*/C*_0_ diverges at a critical pitch of about *P/*(2*πL*) = 0.32. The switch in sign of spontaneous curvature corresponds to an energy barrier in bending energy of the membrane (Fig. 2g).

These results, consistent with our previous results obtained for catenoids of varying neck radius [25], suggest that at least two distinct curvature-inducing mechanisms are required to admit helicoids below the critical pitch as energy minimizers and solutions to Eq. 9. Additionally, the existence of an energy barrier at a specific pitch can be interpreted as a possible biophysical mechanism preventing the collapse of a helicoidal membrane below a critical pitch, and therefore providing a geometrical control of the membrane structure through the Gaussian curvature.

### Ratio between inner and outer ramp radius constrains the distribution of spontaneous curvature in hollow helicoids

Helicoidal ramps observed in organelles are often hollow with a single arm [1, 4, 26] (see Fig. 1a-c). Such a geometry can be constructed by varying *r* ∈ [*r*_0_, *L/*2] in Eq. 7, resulting in a hollow helicoid as depicted in Fig. 3a. We next investigated the relationship between the distribution of spontaneous curvature and the geometry of a hollow helicoid. The presence of the inner ramp radius at *r* = *r*_0_ provides a new geometrical parameter relevant to ER ramp shapes. A natural question that therefore arises is, how does the distance between the inner and outer ramp influences spontaneous curvature in hollow helicoidal membranes? To answer this question, we first impose uniform boundary conditions in spontaneous curvature (*C* = *C*_0_) at the outer edge of the helicoid, while imposing no-flux conditions at the other boundaries. We solve Eq. 9 for the spontaneous curvature on the helicoid surface for fixed pitch (*P/*2*πL* = 0.05), and varying ratio between inner ramp radius and helicoid diameter (*r*_0_*/L*).

**Figure 3:**
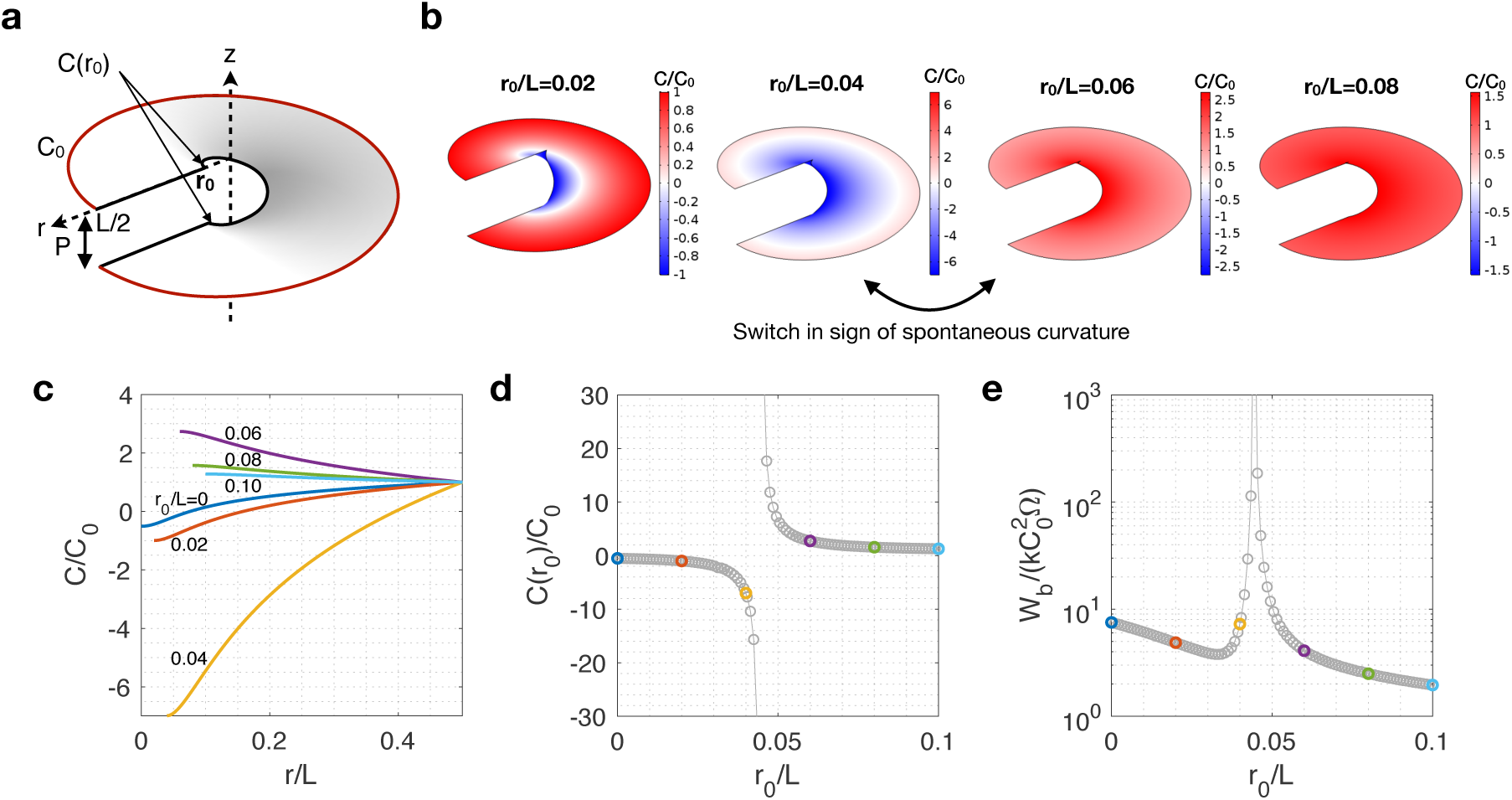
Hollow helicoidal ramps show a switch in the sign of spontaneous curvature that corresponds to an energy barrier at the inner radius when subject to constant exterior spontaneous curvature. (a) Geometry of a hollow helicoid with constant spontaneous curvature *C*_0_ at the outer boundary. (b) Distribution of spontaneous curvature on the surface of hollow helicoids with various inner radii show a sign switch of spontaneous curvature at the center between *r*_0_*/L* = 0.04 and 0.06. (c) Distribution of spontaneous curvature along the helicoid radius for various inner ramp radii. (d) Spontaneous curvature at the inner boundary as a function of the inner ramp radius shows a discontinuity at *r*_0_*/L* ≃ 0.045. (e) Normalized bending energy of the helicoid shows an energy barrier at the critical radius of the inner hole. Color symbols in (d) and (e) match plots in (c). All computations are for helicoids with a pitch of *P/*(2*πL*) = 0.05.

The resulting distribution of spontaneous curvature is shown as a color map on the helicoid surfaces in Fig. 3b, and quantitatively along the helicoid radius in Fig. 3c. We find that for holes with large radii, the spontaneous curvature required to maintain the hollow helicoid is higher at the center, and remains of the same sign as *C*_0_. However for small hole radii, *C/C*_0_ is negative at the center, indicating the presence of a switch in sign at a critical inner radius. This switch, is better visualized when plotting *C/C*_0_ at the inner boundary as a function of the inner radius (Fig. 3d), corresponds to a barrier in bending energy in the membrane (Fig. 3e). A local minimum in bending energy is obtained for values of the ratio between the inner and outer helicoid ramps smaller than for the barrier. Note that, although qualitatively similar to the results obtained for the full helicoid (Fig. 2), the position of the switch and the energy profile have different values.

Next, we explored how the pitch of the helicoid modulates the switch found in hollow helicoids and the associated energy barrier (see Figs. S1 and 4 respectively). As seen from Fig. 4, decreasing the helicoid pitch lowers the value of the radius ratio at which the energy barrier and local minimum occur. Additionally, the local minimum is deeper and narrower. For completeness, we investigate how imposing a constant spontaneous curvature at the inner boundary – instead of the outer – influences our findings. The resulting radial distribution and total bending energy are shown in Fig. S2 and S3 respectively for various hole radii and pitches. No switch-like behavior is observed for spontaneous curvature imposed at the inner boundary. We find that the intensity of spontaneous curvature and bending energy increases with decreasing hole radii and increasing pitch. It should be noted that, by correspondence with catenoids, for any configuration where *r*_0_ > 0, no switch in the sign of the spontaneous curvature occurs if spontaneous curvature is imposed at both inner and outer boundary [25].

**Figure 4:**
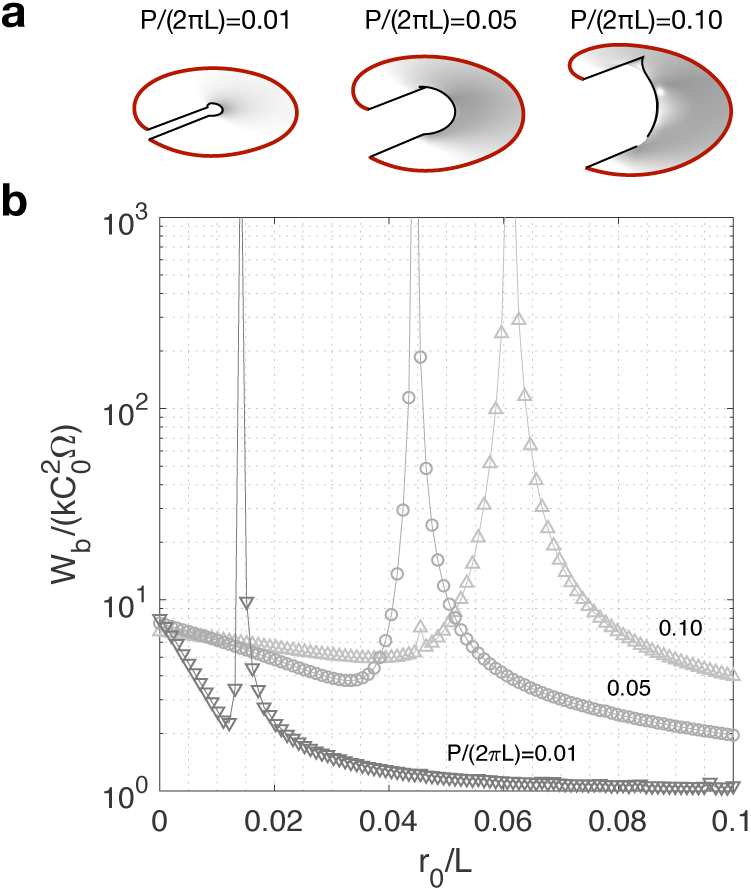
The pitch of a hollow helicoid controls the value of the critical inner radius and the width of the corresponding energy barrier. Helicoids with small pitch exhibit a deeper local energy minimum. (a) Helicoids with three different pitches. (b) Normalized bending energy of helicoids with pitches corresponding to (a), as a function of the inner radius

### Axial gradient of spontaneous curvature at the boundary induces two switches resulting in a locally favorable helicoidal geometry

Next, we investigate how a differential in curvature-inducing proteins along the helicoid axis affects the system. Such configuration is representative of possible inhomogeneous regions of protein source or consumption within the intracellular volume. We thus impose a linear gradient of spontaneous curvature ranging from *C*_0_ to *C*_1_ at the exterior boundary (Fig. 5a). Surprisingly, we find that two distinct switches in spontaneous curvature sign exist at different critical inner radii. These switches correspond to two different limits of *C* at the top and bottom points of the inner ramp and they behave differently for the two critical radii (Fig. 5b,c). At the largest critical radius, the spontaneous curvatures at the top and bottom of the inner ramp both tend to positive infinity for values of the inner ramp radius approaching the critical value from above, and tend to negative infinity when approaching from below. For simplicity, we call this type of switch in spontaneous curvature sign an “aligned switch”. At the smallest critical radius, however, the spontaneous curvatures at the top and bottom ramps tend to opposite limits when approaching the critical value. We therefore call this behavior an “anti-aligned switch”. Both switches correspond to an energy barrier, therefore creating a new local minimum bound by virtually infinite energy barriers (Fig. 5d). As a consequence, if the exterior boundaries are subject to a gradient of spontaneous curvature such as one studied here, then a helicoidal membrane whose inner radius is constrained in-between the two energy barriers will tend to be energetically stable with respect to variations in the inner ramp radius.

**Figure 5:**
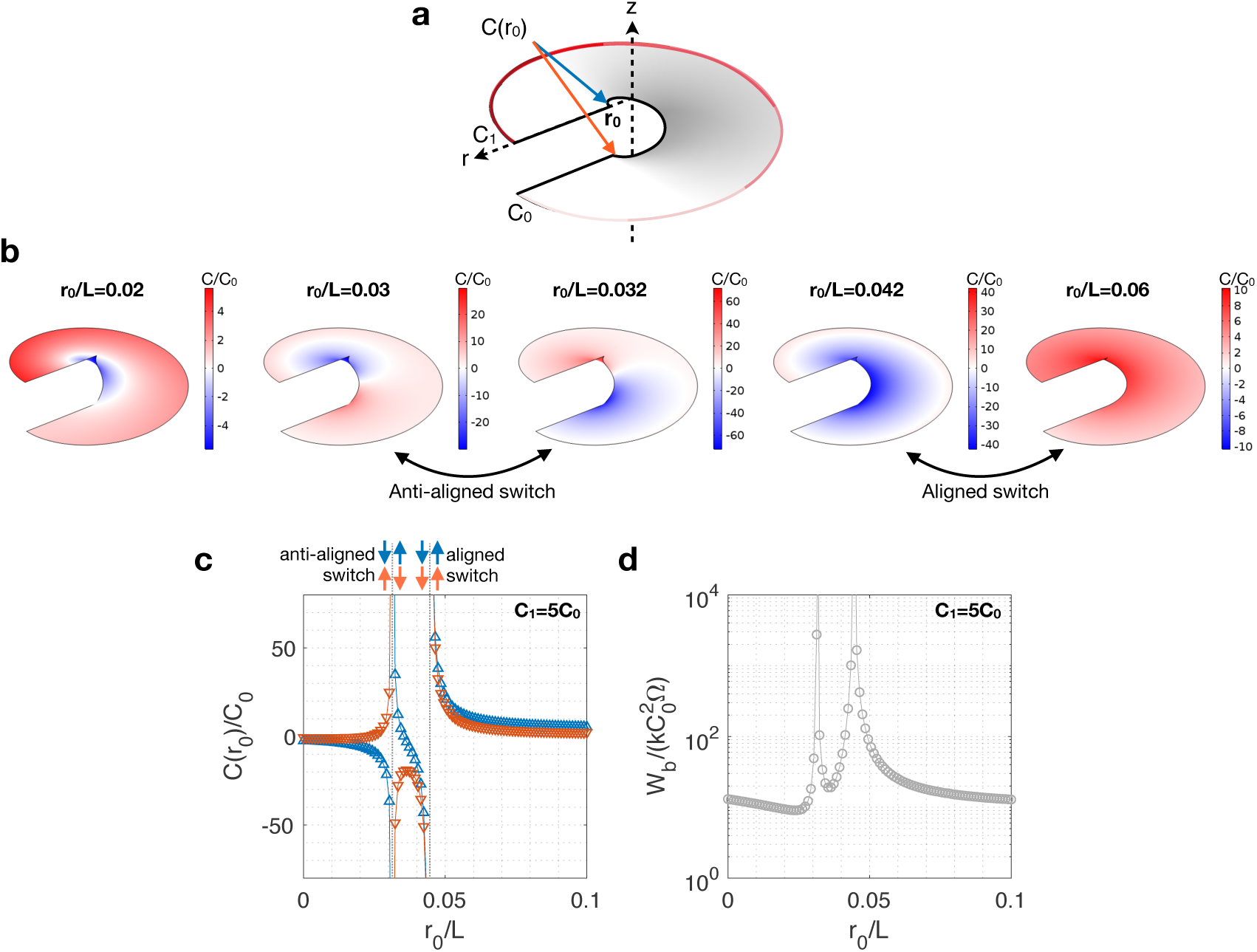
Helicoid with a linear gradient of spontaneous curvature along the external boundary show two distinct sign switches in *C* at the center. (a) Hollow helicoid with an imposed spatial gradient of spontaneous curvature from *C*_0_ to *C*_1_ at the outer boundary. (b) Computed surface distribution of spontaneous curvature on hollow helicoids with *C*_1_ = 5*C*_0_ at various inner radii. Two distinct switches in spontaneous curvature are found at two critical inner ramp radius. All computations are made for helicoids of pitch *P/*(2*πL*) = 0.05. (c) Computed spontaneous curvature at the inner top (upward blue triangles) and inner bottom (downward red triangles) boundary for *C*_1_ = 5*C*_0_. Top and bottom points exhibit two switches in sign of *C* with varying inner radius, however their sign is opposite at the first switch (anti-aligned switch), but equal at the second switch (aligned switch). (d) Each switch corresponds to an energy barrier. The local energy minimum between the two energy barrier represents a possible mechanism for regulating the inner radius of an ER helicoidal ramp through spontaneous curvature.

The existence of two energy barriers corresponding to different switch mechanisms in spontaneous curvature leads to new questions: How is the two-switch behavior controlled by the boundary gradient? How can a membrane structure access the state of local energy minimum between two energy barriers? How does the system transition from a single energy barrier for homogeneous boundaries (Figs. 3 and 4) to two barriers when a gradient in spontaneous curvature at the boundaries is imposed? To answer these questions, we next investigated how the aligned and anti-aligned switches are controlled by the spontaneous curvature gradient at the exterior boundary. To do so, we compute the spontaneous curvature at the top and bottom of the inner ramp for various *C*_1_*/C*_0_ ratios and fixed helicoid pitch (*P/*(2*πL*) = 0.05). As shown in Fig. 6, the aligned and anti-aligned switches vanish at a distinct value of the boundary gradient in spontaneous curvature. This mechanism can be understood using simple symmetry arguments: for *C*_1_ = *C*_0_ no variation in spontaneous curvature in the z-direction can exist, implying that the anti-aligned switch must vanish (Fig. 5 and Fig. 6f). On the contrary, for *C*_1_ = −*C*_0_, the boundary condition imposes the spontaneous curvature along the z-axis to be symmetric with respect to the mid-plane of the helicoid. This therefore requires the aligned switch to vanish (Fig. 6b). Every time the inner ramp radius passes a critical value, the limits in spontaneous curvature associated with the vanishing switch take opposite signs.

**Figure 6:**
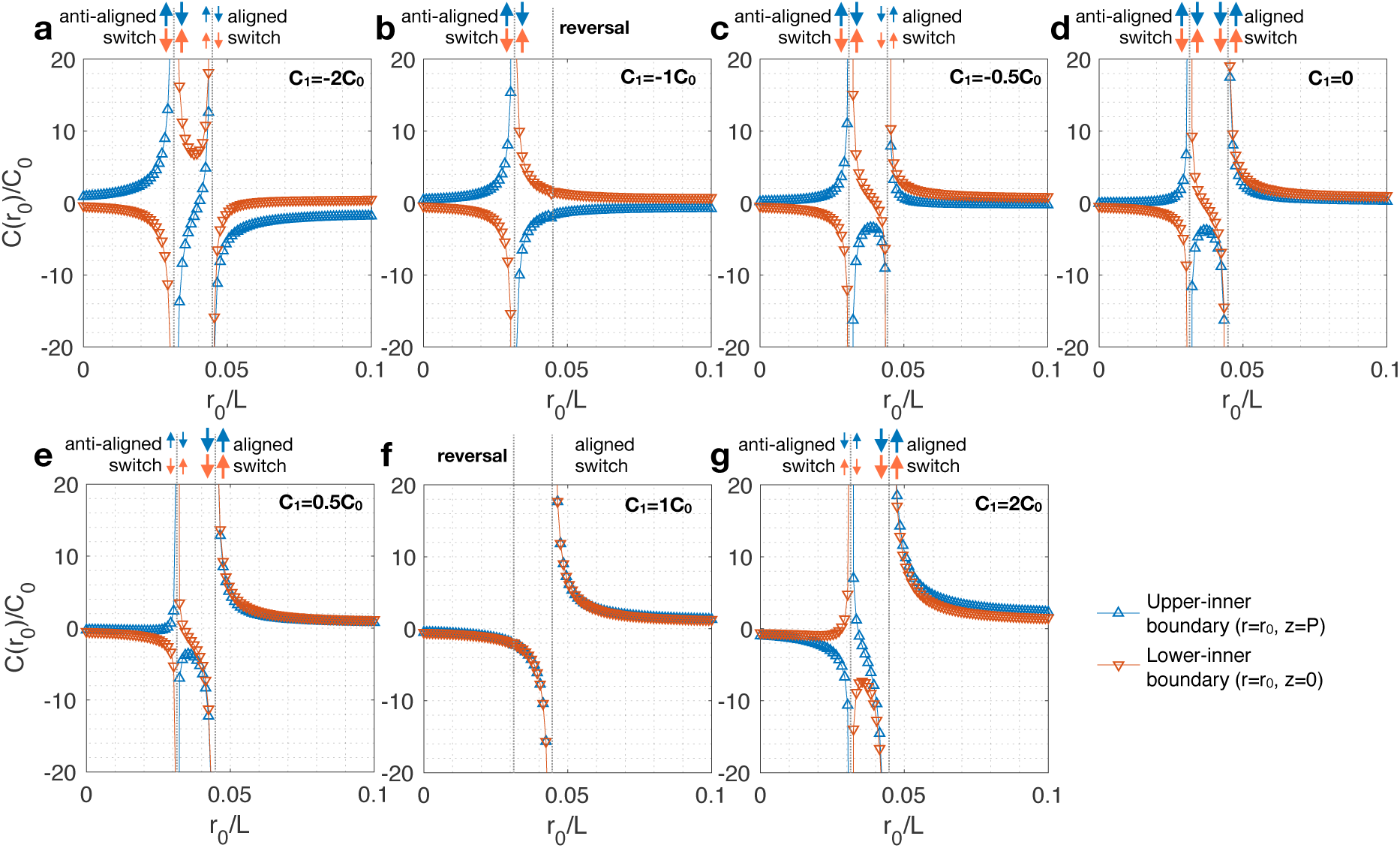
The two switches induced by a gradient in spontaneous curvature reverse their sign at distinct values of the gradient. (a-g) Spontaneous curvature at the upper and lower inner boundary as a function of the inner radius for various gradients at the exterior boundary. (f) For *C*_0_ = *C*_1_, the distribution of C must be independent of *z*, requiring the suppression of the anti-aligned switch. (b) For *C*_1_ = *C*_0_, symmetry requirements induce the aligned switch to be suppressed.

The bending energy for each of the configurations examined in Fig. 6 are shown in Fig. S4, where it can be verified that the energy barrier associated with the switches indeed vanish at the corresponding spontaneous curvature gradients. Finally, we examine the effect of the helicoid pitch on the system. We find that, similarly to the case where the boundary condition is homogeneous (Fig. 4), larger helicoid pitches lead to larger critical values in inner ramp radius, and higher energy values (Fig. S5).

## Discussion

In this study, we proposed a minimal model for computing the spatial distribution of spontaneous curvature on helicoidal structures representative of ER ramps [1, 4, 26]. We identified conditions where continuous variations of the helicoid pitch and inner ramp radius are associated with discontinuous switches in the sign of spontaneous curvature, demonstrating a unique coupling between Gaussian curvature and protein-induced curvature. Membrane shapes associated with these switches correspond to energetically unfavorable configurations, resulting in energy barriers in the corresponding geometric parameter space. Surprisingly, we find that when a gradient of spontaneous curvature is imposed on the external ramp of hollow helicoids, the variation of the inner ramp radius is associated with two distinct switches in curvature sign and energy barriers. Our results suggest a geometric state-space for helicoidal membranes in which a well-constrained energetically favorable state can be accessed by turning-off one or the other energy barrier independently through curvature distribution.

In the context of biological membrane structures, the presence of a switch in sign of spontaneous curvature suggests the requirement for at least two distinct curvature-generating mechanisms to modulate the geometry. These can be thought of as either two different kind of membrane-bound proteins inducing spontaneous curvature of opposite sign, or as the same curvature-inducing protein but inserted on the opposite sides of the surface [33, 40]. Contrary to catenoid-shaped necks where such two-step molecular mechanisms have been identified in mammalian cells (e.g. ESCRT mediated budding [10, 51]) and yeast (e.g. mitochondrial fusion by synergetic action of Dnm1 and Fis1 [27]), the particular mechanisms that relate quantitative structural analysis to protein distribution and activity are not yet available for helicoidal ER geometries. However, a number of curvature-inducing proteins have been suggested to play a role in certain specific ER ramps, including reticulons and DP1/REEP5/Yop1 family proteins in highly curved membrane regions such as tube and cisternae edges [47, 52, 53].

The exquisite coupling between membrane geometry and distribution of curvature-inducing proteins in helicoids has several potential implications in the regulation of ER ramp shapes. The energy barrier associated with pitch and inner ramp radius might help prevent the collapse of ramps below critical geometrical parameter values. Additionally, in the case where a gradient of spontaneous curvature is imposed at the boundaries (Figs. 5c and S4), the helicoid geometry bound by the two energy barriers is energetically stable with respect to variations in the inner ramp radius. Therefore, inducing a gradient of curvature-inducing proteins along the vertical axis of a helicoidal ramp might serve as a possible molecular mechanism to regulate its geometry. Finally, from the reverse perspective, geometric control of the helicoid could present a mechanism to regulate the distribution of proteins on the ER surface. For instance, several proteins are found on the surface of ER cisternae, including Climp-63, kinectin, and p180 [47]. These proteins have transmembrane domains linked to coiled-coil structures on either side of the membrane, and interact with microtubules, ribosomes, or kinesins, and may induce curvature [47, 53]. The relationship between ER shape and distribution of spontaneous curvature identified in this study might underly spatial regulation mechanisms of surface proteins on helicoidal sheets.

Our ongoing work on spontaneous curvature distribution in minimal surfaces has implications for other membrane surfaces as well. In particular, triply periodic minimal surfaces (TPMS) have been observed in various membrane-bound organelles in stressed, diseased, and viral infected cells [54–58], as well as in reconstituted lipid membrane systems [12, 59–63]. In the context of the ER, cubic TPMS have been found in response to elevated levels of specific membrane resident proteins such as cytochrome b(5) [56]. When applied to P-Schwartz and D-Schwartz TPMS with prescribed surface average of spontaneous curvature per unit cell [64, 65], simulation results from our model for minimal surfaces showed that the dimensionless spontaneous curvature qualitatively follows the Gaussian curvature (Fig. 7, see Supplementary Material Section S3 for details [66]). Indeed, *C/C*_0_ is smaller on “flatter” surfaces and larger on the more pronounced saddle-like surfaces.

**Figure 7:**
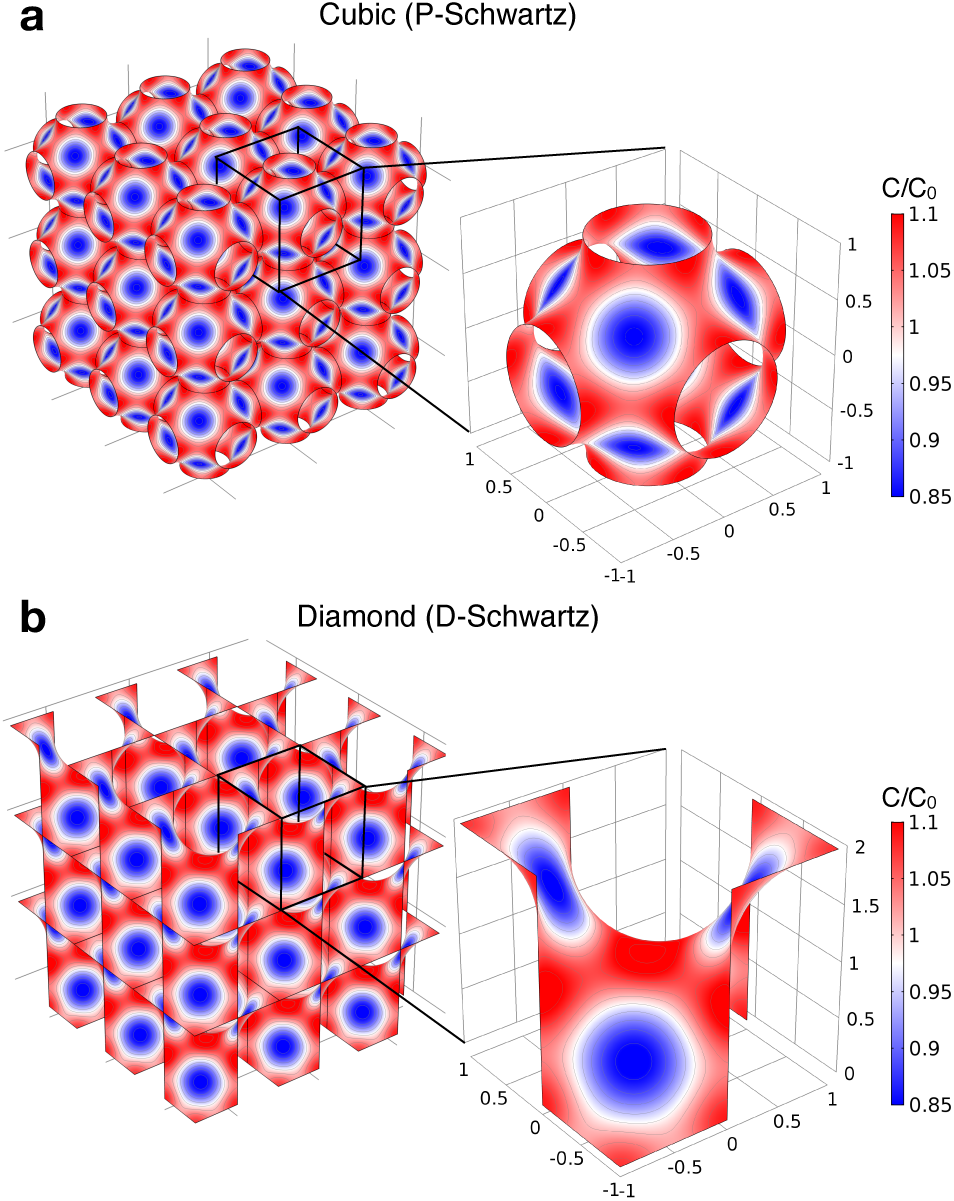
Distribution of spontaneous curvature maintaining a triply periodic minimal surface (TPMS) structure for an imposed surface average of spontaneous curvature of *C*_0_. (a) Cubic P-Schwartz, and (b) diamond D-Schwartz, shown in 3 by 3 arrays and in close up unit cells. Because P- and D-Schwartz surfaces belong to the same associated family of surfaces, the distribution of spontaneous curvature also follows an isometric transformation between the two structures.

In this study we considered helicoidal surfaces as a simplified model for ER ramps observed in the peripheral ER of neural and secretory cells [1, 26]. This simplification results in certain limitations. First, ER sheet structures are made of two lipid bilayer typically 40 nm apart [26, 52]. In the current model, the double bilayer is represented as a unique surface of zero thickness lying in between the two bilayer (see Model assumption (a)), in line with previous modeling studies [26, 30]. This approximation has two consequences on the interpretation of our results in the context of ER structures. First, the effective bending and Gaussian moduli have twice the value of a single bilayer, provided that both bilayers have the same spontaneous curvature [67]. This does not affect the results presented here because on minimal surfaces both moduli vanish from the shape equation and incompressibility condition, and only their ratio appears in the definition of the dimensionless bending energy (Eq. 11). Second, the edges of helicoidal ER ramps are bent surfaces connecting the two bilayers. To account for this geometry, previous modeling studies proposed effective boundary conditions that assume toroidal surface elements at the ramps’ edges [26, 30]. The resulting edge energy was then expressed as analogously to the bending energy for the geodesic curvature, where edge scaffolding proteins induce a spontaneous edge curvature [30]. In the absence of available data supporting the detailed structures of ER sheet edges, we assumed for simplicity either Dirichlet or no-flux Neumann boundary conditions for the spontaneous curvature. The derivation of rigorous effective boundary conditions relating protein distribution and spontaneous curvature at open double-bilayer edges will likely require relaxing the Model assumption (f) to account for area difference elasticity between the leaflets as well as between the two bilayers, and possibly allow lipid tilt at the highly curved edges [40].

While in this paper we mostly discuss the role of protein-induced spontaneous curvature, it is worth noting that asymmetric lipid composition between the inner and outer leaflets can also lead to spontaneous curvature effects (see Model assumption (b)). Depending on the ratio between the size of the lipid headgroup and the surface area occupied by their hydrocarbon chain, lipid shape can be divided between cylindrical, conical, and inverted conical. The two predominantly found lipids in mammalian ER membranes are the cylindrical phosphatidylcholine (PC), and the conical phosphatidylethanolamine (PE) [68]. As a consequence, an unbalanced PC/PE composition between these two lipids across the ER bilayer can generate spontaneous curvature, while an imposed membrane curvature can lead to lipid sorting to accommodate the constrained geometry [69, 70]. In addition to lipid packing effects, membrane electrostatics can also contribute to curvature generation. Indeed, in early endosomal membranes such as in the ER, phosphatidylserine (PS) – a negatively charged lipid primary responsible for membrane electrostatics in cells – is found almost exclusively in the luminal leaflet[71, 72]. It should be noted however that in the present model the surface represents two bilayers. Therefore, lipid composition asymmetry between the luminal and cytosolic sides of the ER are likely to be compensated by the symmetry of the two bilayers with respect to the mid-lumen plan.

Another key assumption of our model is the homogeneous values of bending and Gaussian moduli (Model assumption (g)). One can expect that a heterogeneous membrane composition – either in proteins or lipids – will induce spatial variations of the membrane mechanical properties [73]. The shape equation for heterogeneous membranes only has extra terms proportional to the covariant and surface derivatives of the Gaussian modulus [38, 42]. This means that in the case where only the bending modulus is dependent on the local composition (*k*(*σ*) and *k*_*G*_ is a constant), Eq. (4) remains valid at the condition that spontaneous curvature is reinterpreted as the quantity *C*\*(*σ*) = *k*(*σ*)*C*(*σ*). Additionally, it can be shown that in this case Eq. (6) remains unchanged, and therefore our results hold with *C* = *C*\*. Given the difficulties to measure and compute the value of the Gaussian modulus, such a scenario is a reasonable assumptions. In the case where the Gaussian modulus is also heterogeneous, extra coupling terms appear in the shape and incompressibility equations for the membrane, and a more general model would be required.

Overall this work provides a modeling framework to gain information on the distribution of curvature-inducing proteins on cellular membrane based on the knowledge of their shape. Despite its simplicity, we anticipate that the proposed approach has the potential to support super-resolution microscopy techniques by relating observed membrane structure to their biophysical properties.

## Author Contributions

All authors contributed to the study design, model development, result analysis, and writing of the manuscript. MC carried the numerical implementation and data analysis.

## Acknowledgments

The authors are grateful to Prof. Andreas Carlson, Jennifer Fromm, and Allen Leung for their critical comments.

## Data Accessibility

The main Comsol file, Matlab code, and a sample data file, are available at the author’s Github depository:

github.com/mchabanon/helicoid_spontaneous_curvature

## Funding Statement

This work was supported by the ONR N00014-17-1-2628 and NSF PHY 1505017 awards to PR.

## Supplementary Material

### S1 Local force balance of elastic surfaces at mechanical equilibrium

The equation of mechanical equilibrium of an elastic surface *ω* subject to a lateral pressure *p* can be written in the compact form

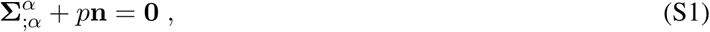

where **Σ**^*α*^ are the stress vectors and **n** is the unit normal to the local surface. Greek indices range over 1, 2, and if repeated, are summed over this range. Semicolon identifies covariant differentiation with respect to the surface metric *a*_*αβ*_ = **a**_*α*_ · **a**_*β*_ where **a**_*α*_ = **r**_,*α*_ are the tangent vectors and **r**(*θ*^*α*^) is the parametrization of the position field. The commas refer to partial derivatives with respect to the surface coordinates *θ*^*α*^. With these definitions, the normal vector is given by **n** = (**a**_1_ × **a**_2_)*/* | **a**_1_ × **a**_2_ |. In Eq. S1, the differential operation represents the surface divergence defined as 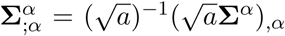 where *a* = det(*a*_*αβ*_). In surface theory, a manifold is described by the metric *a*_*αβ*_ defined above, and the curvature tensor given by *b*_*αβ*_ = **n** · **r**_,*αβ*_.

For an elastic membrane whose energy surface density per unit mass depends on the metric and curvature only *F* (*a*_*αβ*_, *b*_*αβ*_; *θ*^*α*^), the stress vectors involved in the local force balance (Eq. S1) can be written as [42]

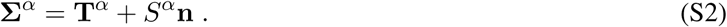

Here the tangential stress vectors are

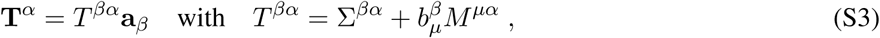

and the components of the normal stress vectors are

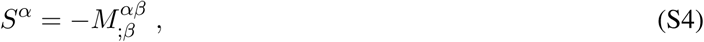

where 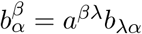. The components of the stress vectors depends on the energy density as [42]

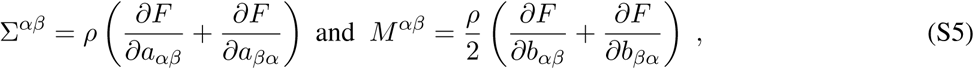

where *ρ* is the surface mass density of the membrane. The tangential and normal local force balances can now be obtained by introducing Eqs. S2, S3, and S4 into Eq. S1, resulting in

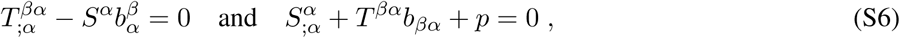

where we made use of the Gauss and Weingarten equations [45] **a**_*α*;*β*_ = *b*_*αβ*_ **n** and 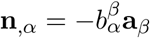 respectively.

The free energy density can be written as a function of the mean curvature *H* and Gaussian curvature *K*. These are related to the metric and curvature by

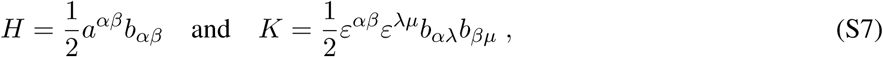

where *a*^*αβ*^ = (*a*_*αβ*_)^−1^ is the dual metric, and *ε*^*αβ*^ is the permutation tensor defined by 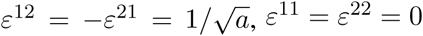. According to this definition (Eq. S7), the free energy density per unit mass can be re-written in terms of the mean and Gaussian curvatures *F* (*H, K*; *θ*^*α*^). Furthermore, lipid membranes are essentially incompressible (see assumption (d) in the Model Development Section of the main text). This is imposed using a Lagrange multiplier *γ*(*θ*^*α*^) to ensure that the local area dilatation *J* = 1, or equivalently, to constraint the constant surface density *ρ* of the membrane. Consequently we can define the surface energy density of the membrane as follows

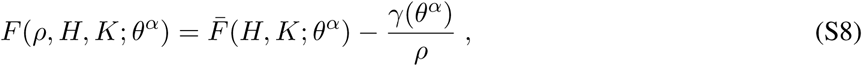

and when introducing the surface energy per unit area 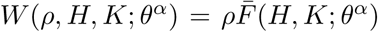, the components of the stress vectors (Eqs. S5) can be written as [42]

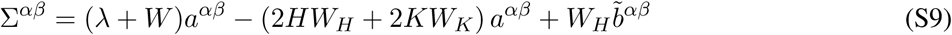

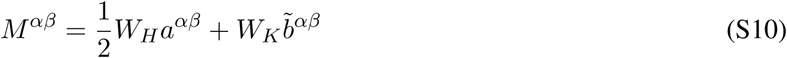

where *λ*(*θ*^*α*^) = −[*γ*(*θ*^*α*^) + *W* (*H, K*; *θ*^*α*^)], and 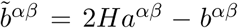 is the cofactor of the curvature. The subscripts *H* and *K* refer to the partial derivative of the energy with respect to the indicated variable. Note that the Lagrange multiplier *γ* can be interpreted as a surface pressure, and is not a material property of the surface [35, 38]. Consequently, *λ* can be interpreted as the surface tension based on comparisons with edge conditions on a flat surface [35].

Finally, introducing Eqs. S9 and S10 into Eqs. S3 and S4, we can rewrite the normal and tangential force balances (Eqs. S6) as

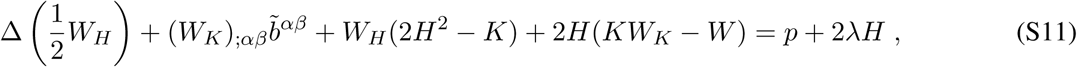

and

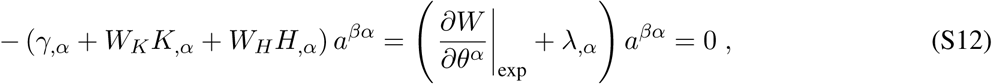

where 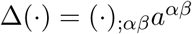 is the surface Laplacian (or Beltrami operator), and *∂*(·)*/∂θ*^*α*^|_exp_ is the explicit derivative with respect to *θ*^*α*^.

Eqs. S11 and S12 are the general shape equation and incompressibility condition for an elastic surface with free energy per unit area *W* (*ρ, H, K*; *θ*^*α*^).

### S2 Helicoid to catenoid transformation

According to the Gauss’ *Theorema Egregium*, the distribution of Gaussian curvature on a minimal surface follows any isometric mapping of such surface [49]. As illustrated in Fig. 2b, such transformation exists between helicoids and catenoids.

**Table S1:** Command sequence to refine the surface meshes of the elementary P- and D-Schwartz elementary surfaces. Description of the commands can be found in the Surface Evolver documentation.

| Elementary surface | Surface evolution sequence |
| --- | --- |
| P-Schwartz | g 5; r; g 5; r; g 5; r; g 10; u; r; g 10; u; r; g 10; u; u; refine edge where on_constraint 2; refine edge where on_constraint 4; hessian; hessian; u ; |
| D-Schwartz | g 5; r; g 5; r; g 5; r; g 10; u; r; g 10; u; r; g 10; u; u; hessian; hessian; r ;<br>g 100; hessian; hessian ; g 100; u; g 500 ; hessian; g 100; |

Helicoids and catenoids belong to the same associated family of surfaces. The (continuous) isometric transformation from one surface to the other can be written as

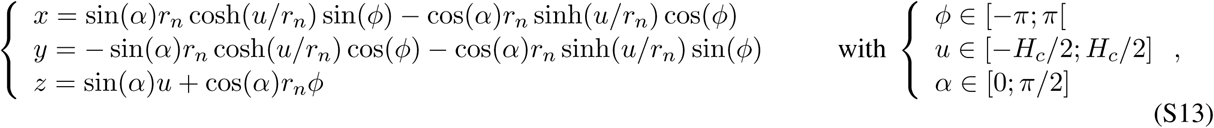

where *P* is the helicoid pitch, related to the catenoid neck radius by *r*_*n*_ = *P/*(2*π*), and *L* is the helicoid diameter, related to the catenoid heigh by *H*_*c*_ = 2*r*_*n*_ sinh^−1^(*L/*(2*r*_*n*_)).

For *α* = *π/*2, this system describes a catenoid such as the one shown in Fig. 2c, while for *α* = 0 it describes a helicoid as in Fig. 2a. The equivalence between Eqs. S13 and 7 when *α* = 0 can be verified with the change of variable *r* = *r*_*n*_ sinh(*u/r*_*n*_). Any intermediate values of *α* gives rise to a minimal surfaces belonging to the associated family of helicoids and catenoids, as illustrated in Fig. 2b.

The Gaussian curvature of these surfaces is given by

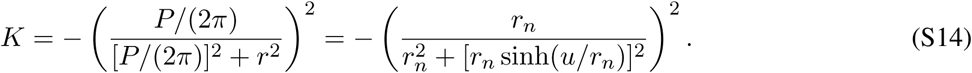

### S3 Method for computing the distribution of spontaneous curvature on TPMS

Elementary surfaces P-Schwartz or D-Schwartz were downloaded from ^1^ and imported in Surface Evolver [66]. The surface energy was minimized through a succession of minimization and mesh refinement operations (see Table S1). The refined elementary surfaces were then duplicated and flipped to produce the periodic cubic unit cells, before to be exported in .stl format ^2^.

The cubic unit cell meshes were imported in Comosol Multiphysics, and Eq. 9 was solved using the “Surface reaction” module. Because of the periodic nature of the TPMS, instead of imposing Dirichlet boundary conditions at the boundaries, we chose periodic flux boundary conditions at the opposite edges, and imposed a surface average of *C*_0_ on the unit cells. This choice is inspired from approaches to solve closure problems in transport in porous media, where the elementary unit volume is assumed to be periodic [64, 65].

### S4 Supplementary Figures

**Figure S1:**
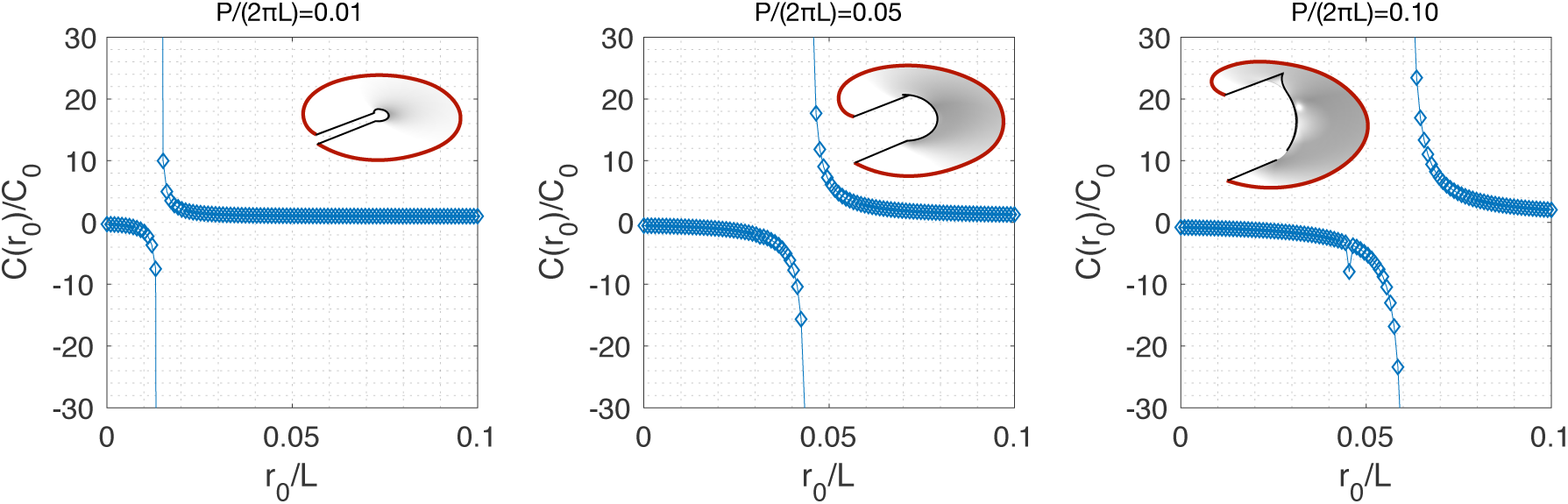
Helicoid pitch controls the position of the energy the spontaneous curvature switch in sign at the inner boundary.

**Figure S2:**
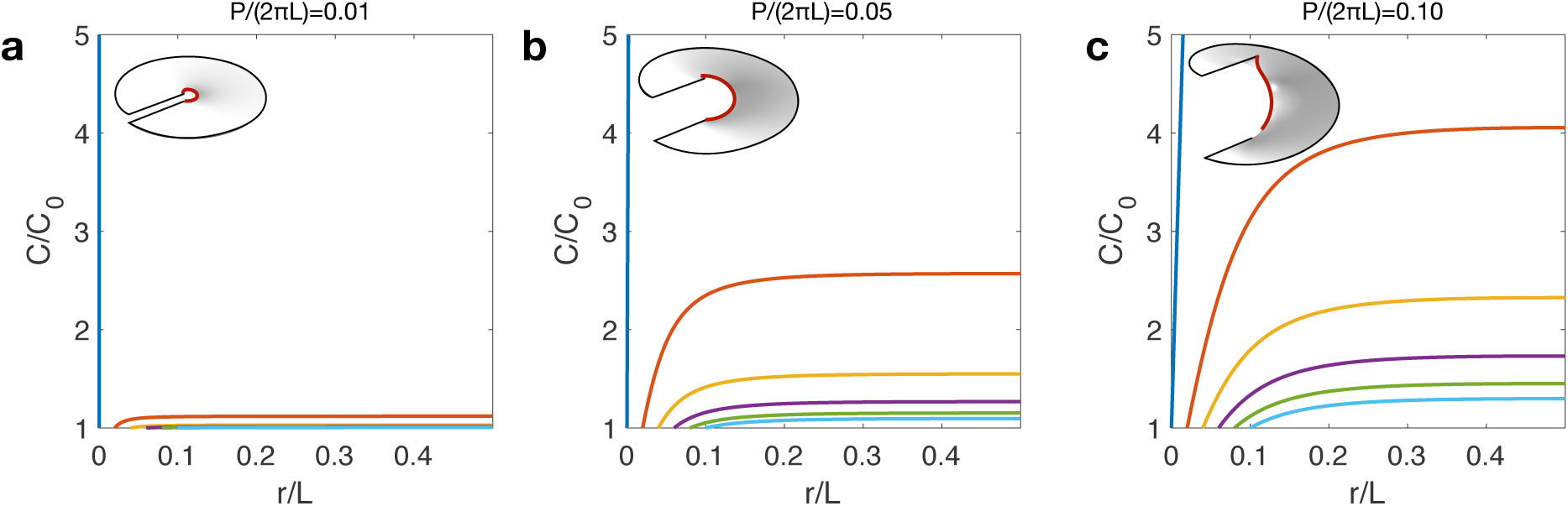
Distribution of spontaneous cutvature along the helicoid radius for *C*_0_ imposed at the inner boundary of the catenoid for various pitch and inner radii. No switch is observed.

**Figure S3:**
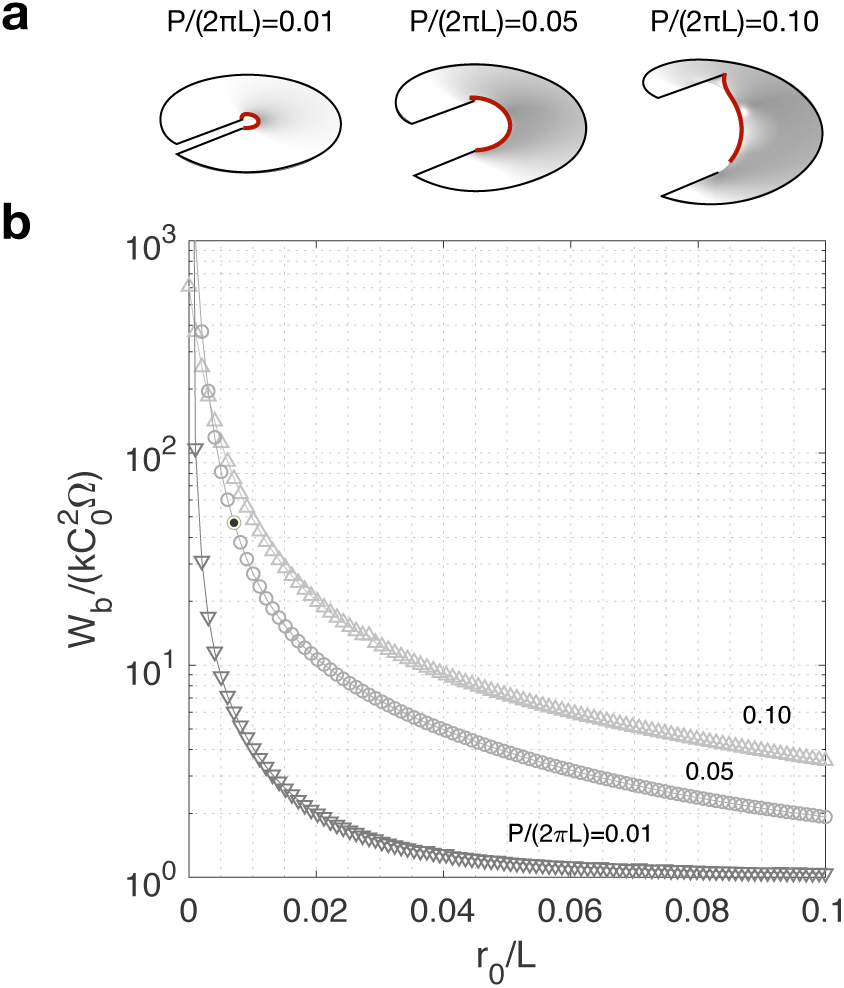
Normalized bending energy of helicoids with boundary conditions imposed on the inner bondary. No energy barrier is observed.

**Figure S4:**
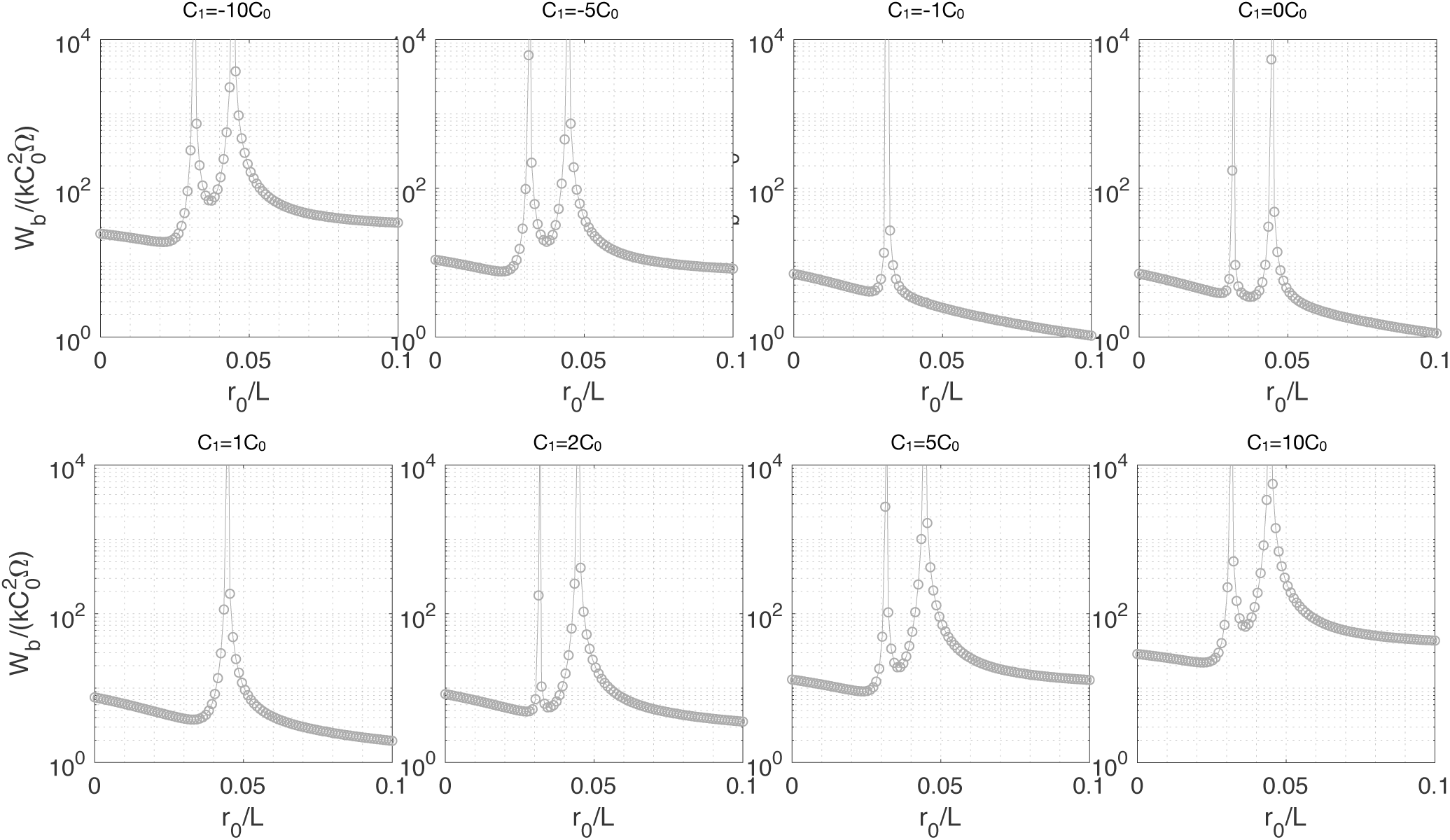
Effect of the imposed gradient in C at the external boundary on the bending energy of the helicoid.

**Figure S5:**
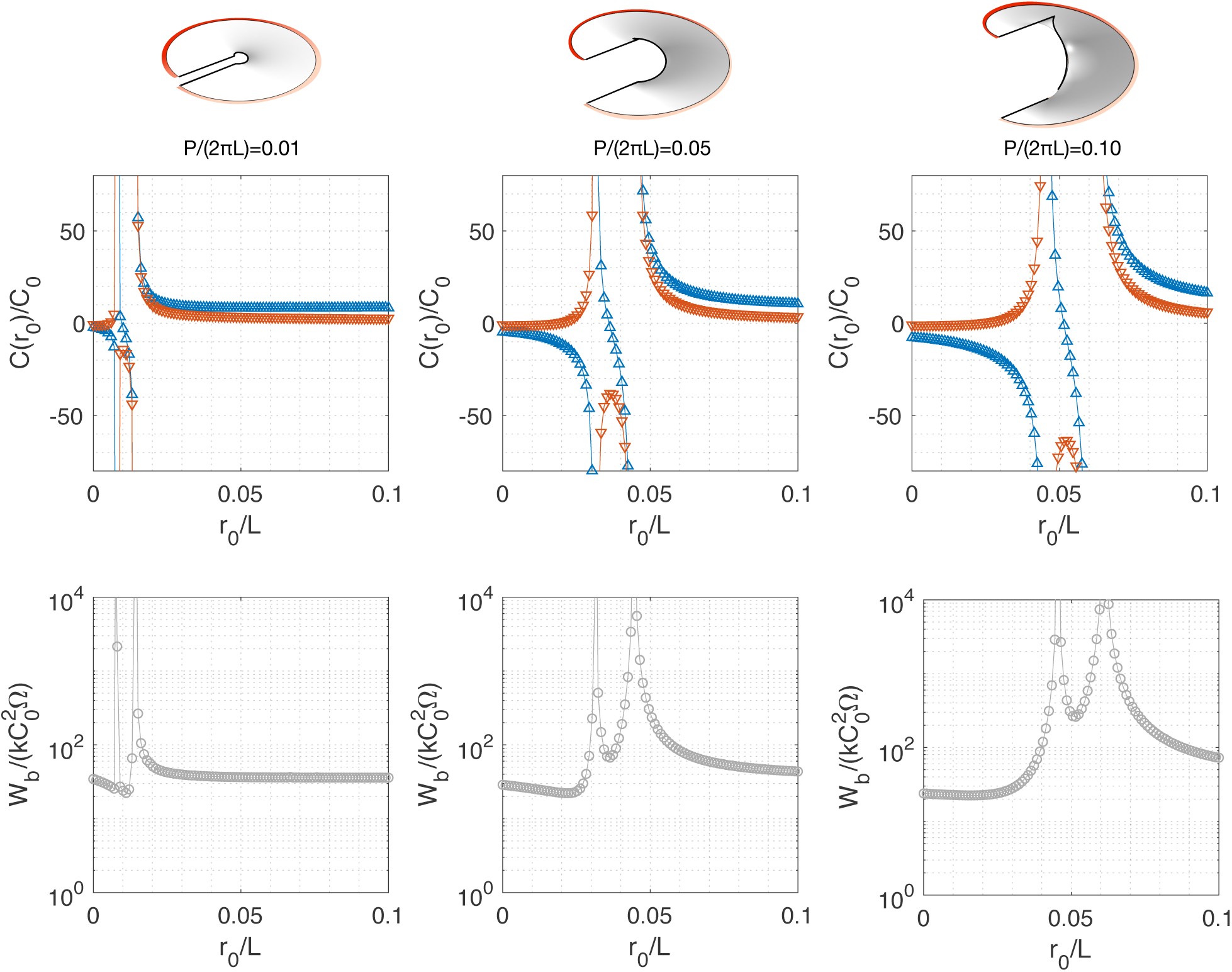
Effect of the pitch on the switch and energy barrier of helicoids with imposed gradient of boundary conditions at the exterior boundary.

## Footnotes

1 facstaff.susqu.edu/brakke/evolver/examples/periodic/periodic.html (consulted on January 29th, 2019)

2 github.com/kashif/evolver/blob/master/fe/stl.cmd (consulted on January 29th, 2019)

## Notes

https://github.com/mchabanon/helicoid_spontaneous_curvature

